# Coherent Optical Scattering and Interferometry (COSI) for label-free multiparameter quantitative imaging of intra-thrombus stability *in vitro*

**DOI:** 10.1101/2020.08.17.254292

**Authors:** Y. Zheng, S. J. Montague, Y. J. Lim, T. Xu, T. Xu, E. E. Gardiner, W. M. Lee

## Abstract

Although existing microfluidics *in vitro* assays recapitulate blood vessel microenvironment using surface-immobilized agonists under biofluidic flows, these assays do not quantify intra-thrombus mass and activities of adhesive platelets at agonist margin and uses fluorescence labeling, therefore limiting clinical translation potential. Here, we describe a real time label-free *in vitro* quantitative imaging flow assay called **C**oherent **O**ptical **S**cattering and phase **I**nterferometry (COSI) that evaluates both intra-thrombus and adhesive-only platelet dynamics using only changes in refractive index. By combining coherent optical scattering and optical interferometry, we evaluated and quantified both intra-thrombus mass with picogram accuracy and adhesive platelet-only events/dynamics with high spatial-temporal resolution (400 nm/s) under fluid shear stress using only changes in refractive index. Using oblique illumination, COSI provide a ∼ 4 µm thin axial slice that quantifies the magnitude of physical of surface adhesive platelets (spreading, adhesion and consolidation) in a developing thrombus without labelling under fluid shear stress. We achieve real time visualization of recruitment of single platelet into thrombus and further correlate it to the developing mass of a thrombus. The adhesive platelet activity exhibit stabilized surface activity of around 2 µm/s and intra-thrombus mass exchange were balanced at around 1 picogram after treatment of a broad range metalloproteinase inhibitor (250 µM GM6001).

**Significance:** The combination of phase imaging with transmitted light and backscattering imaging via oblique illumination in COSI unpicked intra-thrombus mass and adhesive platelet-only activity events at picogram and sub-micrometer precision with millisecond time resolution under fluid shear stress. COSI maps the longitudinal time dynamics of adhesive platelets along changing thrombus mass under metalloproteinase inhibition, and demonstrates potential for real-time correlative microfluidic label-free imaging for flow-dependent biological adhesive events.

## INTRODUCTION

Understanding interactions at the thrombus/surface interface using microfluidic channels coated with vascular wall constituents will enhance our understanding of thrombus evolution process, including platelet activation, adhesion, aggregation, and thrombus stabilization as illustrated in Fig. 1A. It will also advance the biocompatibility of new materials for intravascular devices (1), development of ‘synthetic’ platelets (2) and improve drug delivery (3). Microfluidic channel-based assays for thrombus measurements *in vitro* typically only quantitate either surface-driven biophysical processes associated with platelet activation and spreading (Fig. 1B) or the growth, propagation and stability of a thrombus formed on an immobilized prothrombotic surface (Fig. 1C), but not both. The net stability of a thrombus under a parabolic velocity flow profile (4-8), as shown in Fig.1D, is dependent on a complex set of biophysical parameters. The ability of a thrombus to remain stable can be defined by two key physical interactions. The first is the physical attachment of platelets to a substrate (immobilized agonist), where adhesive platelets, colored red in Fig.1B, aggregate during the initial phase of thrombus formation (9). The second is the platelet-platelet interactions and thrombus consolidation processes occurring during the accumulation phase, with the net effect represented by changes in thrombus mass, colored in caramel in Fig.1D. Surface-driven effects defined by the activities of basal adhesive platelets, include platelet spreading and formation of podia at the base of a thrombus on an agonist-rich surface. Both the intra-thrombus stability and the surface-driven platelet activities contribute to the overall stability of a thrombus (9); that is, how well thrombi adhere to a substrate-coated surface with minimal changes in mass as it withstands changes to the velocity flow gradients, as the thrombus builds.

**Fig. 1.**
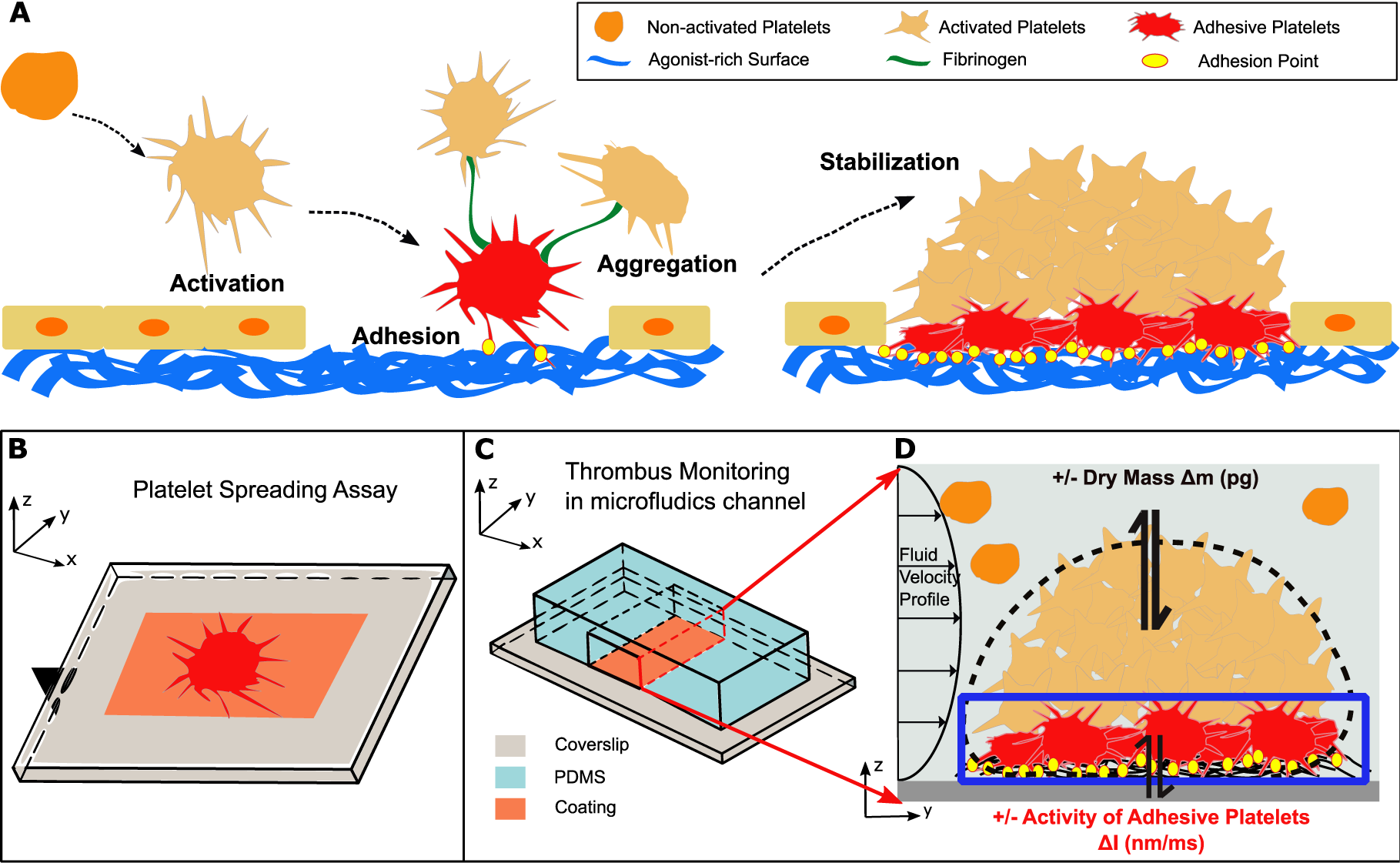
Schematics of changes in mass and local platelet activities in forming and embolizing thrombi. **(A)** The dynamic thrombus formation including four distinct stages: platelet activation, adhesion, aggregation and stabilization. (**B)** The experiment method for studying platelet activation with platelet spreading assays. Cross-section view of a coated microfluidic channel used to visualize surface driven events during platelet spreading. **(C)** The cross-section view of a coated microfluidic channel used to visualize surface driven events during thrombus formation. **(D)** Inside the coated microfluidics channel, a formed thrombus is comprised of platelets interacting with agonist-coated surfaces (collagen/fibrin). The thrombus poses a complex biological system which interacts with the surrounding microenvironment and adapts to changing fluid shear forces. The intra-thrombus stability can be quantified as change in dry mass (colored in caramel) and adhesive platelet activity (colored in red).

In clinical settings, rapid *in vitro* diagnostic tests that quantify intra-thrombus dynamics are frequently conducted without fluorescence labeling. Hemostasis tests, using the platelet function analyzer (10), optical platelet aggregometry (11) and thromboelastography (12), quantify the time to occlude a channel or form aggregates in suspension or on a surface with some physical properties. These clinical tools have several shared limitations, including requirement of a platelet count of at least 100 ⨯10^9^/L, meaning platelet function testing and quantitation of time to reach occlusion cannot be evaluated in people with thrombocytopenia (13, 14). Further, these tests do not inform on changes occurring at a sub-micrometer scale within the thrombus or provide information on the platelet adhesion dynamics during thrombus formation. Thrombus micro-imaging tools that can simultaneously quantify adhesive platelet activity and intra-thrombus mass and that are label-free, do not currently exist, as compared to COSI methods (Table 1).

Platelets are active blood cells without a nucleus and have a relatively stable refractive index. Therefore label-free platelet imaging methods can exploit the refractive index of platelets as an endogenous label for reliable microfluidic imaging. However, current platelet label-free imaging employs either classical interferometry (15) i.e. quantitative phase microscopy (QPM) (7, 16), diffraction phase microscopy (17) optical diffraction tomography (18) or near field light scattering i.e. reflectance interference contrast microscopy (RICM) (19). Furthermore, the angle of incidence from both techniques are either projected in transmission or epi-illumination which make it difficult to selectively image only the adhesive platelets. As such, existing label free imaging modalities either records thrombus formation or just single platelet. For example, existing label-free light scattering imaging techniques, such as RICM (19) or iScat (20, 21), measures total scattered light in an epi-illumination configuration that also include out of focus scattering signals and so cannot readily permit selective imaging of adhesive platelets within a developing thrombus. As the result of limiting to single label-free imaging approaches, there is a loss of valuable spatial-temporal information about dynamic adhesive platelet behaviour at the prothrombotic surface especially during thrombus formation. Recently, a new oblique illumination light scattering imaging method called rotating optical coherent scattering (ROCS) (22, 23) was shown to capture surface-only activities of extended membrane folds in an immune cell with sub-micrometer precision on milliseconds time scale (24). The unique feature of this backscattering imaging method permits selective imaging within a pre-determined depth of focus and so can capture surface driven activities. Depth selection can be adjusted simply by varying the oblique illumination angle and also permits the removal of out of focus scattering signal. Hence, we argue that it is ideal to combine interferometry (QPM) with rotational backscattering imaging to enable the study of how initial adhesive platelets behave on agonist-coated surfaces, including collagen and von Willebrand Factor (VWF), whilst simultaneously quantifying intra-thrombus mass (25). The combinatory label-free imaging (interferometry and scattering) will be able to record the rate and extent of thrombus formation together with intra-thrombus stability and adhesive surface-driven activities. Synchronized multi-parameter label free imaging will therefore better inform us on how physical motility of adhesive platelets and overall morphological of thrombus changes in clinical and research settings, and so expand upon standard assessments of thrombus diagnosis.

We term this new combinatory label-free imaging approach as **C**oherent **O**ptical **S**cattering and **I**nterferometry (**COSI**) that signify real time, label-free, correlative quantitative measurement of intra-thrombus stability (changes in thrombus mass in picogram) (26, 27) and surface-driven platelet activities (adhesive platelets spreading and protrusion) with picogram (pg) and nanometer/millisecond (nm/ms) precision. In addition to unravelling multiparameter information of platelets and thrombus in most transparent microfluidic devices, COSI also removes the limitations of single shot off-axis QPM by conducting refractive index calibration (platelet or red blood cells), where refractive index is constant.

We first perform standardized imaging calibration of COSI for standard single platelet spreading assays involving single platelets, platelet-platelet interactions and hyperactive platelets before conducting imaging trials on *in vitro* thrombus formation under fluid shear stress using collagen- or fibrin-coated cover slides. Once we have established that, we then move onto the evaluation of platelet aggregation and thrombus formation. To quantify adhesive platelet behavior and the dynamic changes of thrombus mass, we adapted an image subtraction algorithm (24) for both the backscattering intensity images and phase images, that quantifies spatial-temporal activity of adhesive platelets adhering to prothrombotic collagen fibers and quantify temporal fluctuation (gain and loss) of thrombus mass over a fixed time interval. The combined spatial temporal imaging of both the surface driven activity of adherent platelets and dry mass changes of a developing thrombus under (patho)physiological shear rates allow us to directly evaluate intrathrombus stability and basal surface activity of the thrombus in the presence or absence of metalloproteinase inhibition (giving conditions of reduced platelet receptor shedding).

## Results

### Coherent Optical Scattering and Interferometry (COSI) imaging platform

The COSI imaging setup (Fig.2A) consists of two cameras that collects transmitted light (red line) and backscattered light (blue line) separately. The light transmitted through the sample accrues an optical phase delay *ϕ*_*t*_ imposed by the sample of different refractive index (Fig. 2B(i)). The transmitted light is combined with a plane reference beam before the camera (red box) to form an interference pattern and the dry mass quantification is retrieved from the resulting interferogram. A rotating oblique illumination, on the other hand, illuminates the sample at a set of pre-determined inclined angles radially around the sample. Instead of accruing phase delay, a sequence of backscattered light emerges because of the refractive index difference between sample (n_s_) and the surrounding medium (n_m_). The integration time of the backscattering signal collection camera (blue box, Fig.2A) is preset so the sample is illuminated sequentially from the full azimuthal 2π angle and the scattering signal under each azimuthal angle is numerically summed to remove the scattering artefacts (Fig. 2C(i)) (22). The microfluidics chamber (a pre-coated straight channel) is placed above the imaging objective as shown in Fig. 2A. The inlet and outlet of the channel are connected to a reservoir containing platelet-rich plasma (PRP) and a syringe pump respectively. The pump draws PRP across the coated channel allowing thrombi to form. The COSI imaging system is compatible with most transparent microfluidics devices that use coverslips (#1 or #1.5 thickness) or a glass slide depending on oblique illumination from the objective lenses. Here, we tuned the oblique illumination angle to only collect light at ∼ 2.64 μm in thickness (Fig. S1D), which is not attainable using conventional label-free imaging reflectance contrast methods. An objective with larger working distance and higher numerical aperture may be selected for thicker glass but could be limited in scattering signal collection. Details of the COSI imaging setup and experimental setup can be found in Methods and signal calibration are shown in Fig. S1A respectively.

**Fig. 2.**
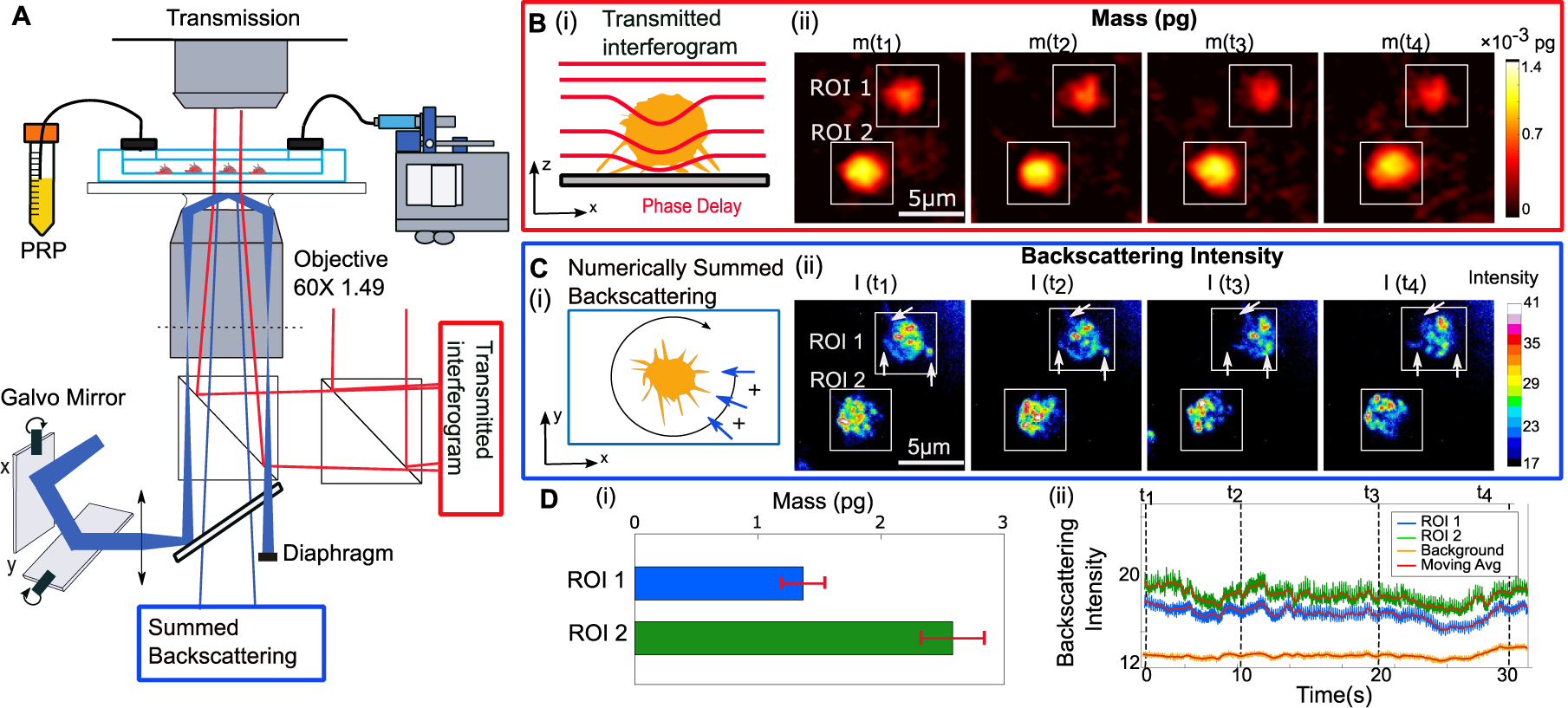
COSI imaging approach to collecting mass distribution and backscattering signal of a sphere. **(A)** Imaging setup of COSI, where red light path represents the interferogram and blue path represents backscattering signal. The mass distribution is retrieved from the phase delay of light transmitted through the sample. The back scattering signal under rotational oblique illumination rejects out of focus light within a variable depth from 200 nm to 4.5 μm. (**B**) (i) The light transmitted through the sample poses a phase delay because of the refractive index difference and this phase delay can be retrieved from the interferogram to quantify the dry mass of the platelet. (ii) Mass distribution map of (***m***(***t***)) of the spreading platelet. (**C**) (i) The sample is illuminated sequentially from the full azimuthal 2π angle and the scattering signal under each azimuthal angle is numerically summed to remove the scattering artefacts. (ii) Back scattering signal (***I***(***t***)) with fluctuating hotspots (arrow). **(D)** (i) Total mass of the selected platelets at different times after spreading. Error bars = Standard Deviation ii) Intensity fluctuation of corresponding ROIs.

### Quantifying platelet spreading, interaction and hyperactivity using COSI

Platelet spreading assays are used routinely to study platelet adhesion and shape change on agonist-coated surfaces, where platelets transform from a rounded, discoidal shape to flattened fully spread platelets, with lamellipodia, filopodia and other membrane protrusions being generated. The resulting measurement of the scattering signals, along the interface with varying refractive index difference indicate that, the maximum intensity is observed along the interface where refractive index is highest. In a platelet, this would mean that the scattering is likely to be observed along large alterations to membrane folding (i.e. ruffles), and where significant refractive index changes along the interface (28, 29). To confirm this, we additionally captured scattering signals from fixed platelets that were stained with phalloidin to visualize actin and compared fluorescence signal with the scattering image (Fig. S2). The scattering images show maximum intensity along the region where actin-rich membrane ruffling and protrusions appear (Fig. S2C and S2D). This is consistent with previous correlative cell migration imaging studies using phase contrast, fluorescence and EM microscopy (29), which indicated a high amount of optical phase delay (destructive interference) occurred along the membrane protrusion that was associated with densely packed bundles of actin filaments. Fig. 2B (ii) and 2C (ii) shows representative COSI imaging frames of two individual platelets within a single imaging field of view, labelled as region of interest (ROI) 1 and ROI 2 <u>(full Movies are available in S1 and S2)</u>. Fig. 2B (ii) and 2C (ii) shows the reconstructed mass distribution (*m*(*t*)) and the backscattering intensity map (*I*_*back*_(*t*)) respectively. COSI was able to distinguish the difference of mass between the 2 platelet; platelet in ROI 1 possess a mass of 1.4 pg while the platelet in ROI 2 had a heavier mass of approximately 2.6 pg (Fig. 2D (i)) that is consistent with a published mean platelet mass measurement of 1.85 ± 0.2 pg (30). The two platelets in ROI1 and ROI2 were shown to have distinctively different physical behavior, display finger like morphologies that are consistent with platelet activation (31) in Fig. 2C(ii)) and very distinct intensity hot spots (200-400 nm in size) that changed within 30 milliseconds (Movie S2). Membrane protrusions were observed to extend and retract at different timepoints (arrows, t_1_, t_3_ and t_4_ (Fig. 2C (ii)). The discrete scattering hotspots of the platelet in ROI 1 displays fewer temporal changes reflecting reduced movement along the fibrin-coated surface. In contrast, the larger platelet (ROI 2) displayed less membrane protrusion but high intensity in hot spots and spatiotemporal movements. The average scattering intensity fluctuations across each platelet for both ROIs indicates greater dynamic local membrane fluctuation for ROI 2 (green) compared with ROI 1 (Fig. 2D (ii)). The results above confirmed that COSI can readily observed changes in dry mass of resting (adherent) and activated (spread) single platelets.

However, a critical drawback in the scattering images only qualitatively shows some fluctuations and does not quantitate magnitude of motility, spreading and interaction.

To further quantify the magnitude of the platelet activity that would be suitable under flow, we implemented an image subtraction method (23, 24) which clearly quantify the magnitude of physical activity of highly activated platelets. The subtraction method works by calculating difference of intensity (Δ*I*(*t*)) at each digital pixel along a 2D matrix between sequential frames (24). For a calibrated 2D array (pixel/nm), the subtraction method quantifies the rate of change of the object in a given time interval. The calculated rate of change can be defined by the magnitude of the activity of platelets. In contrast to existing kymography (32), this method also aids in removing static objects. Fig. 3A (i) illustrates this concept with a simplified diagram using a 2D grid representing individual pixels on a camera screen, and an irregular object with a given scattering profile represented by an intensity I (x,y,t_0_) pattern, colored yellow. The object moves from position x_0_ to x_1_ within a certain time interval Δt from time t_0._ The subtraction of two images, I (t_0_) from I (t_0_-Δt), highlights the leading edge of the object and the direction of its movement along the lateral plane within a fixed depth of focus. Hence the higher the difference of intensity of light scattering (Fig. 3A (ii)) is correlated with an increase in lateral platelet activity that is a direct attribute of membrane ruffling and cytoskeletal rearrangement. The spatiotemporal resolution of the subtraction method is determined by temporal imaging speed of each frame i.e. time interval Δt and pixel resolution of the system (nm).

**Fig. 3.**
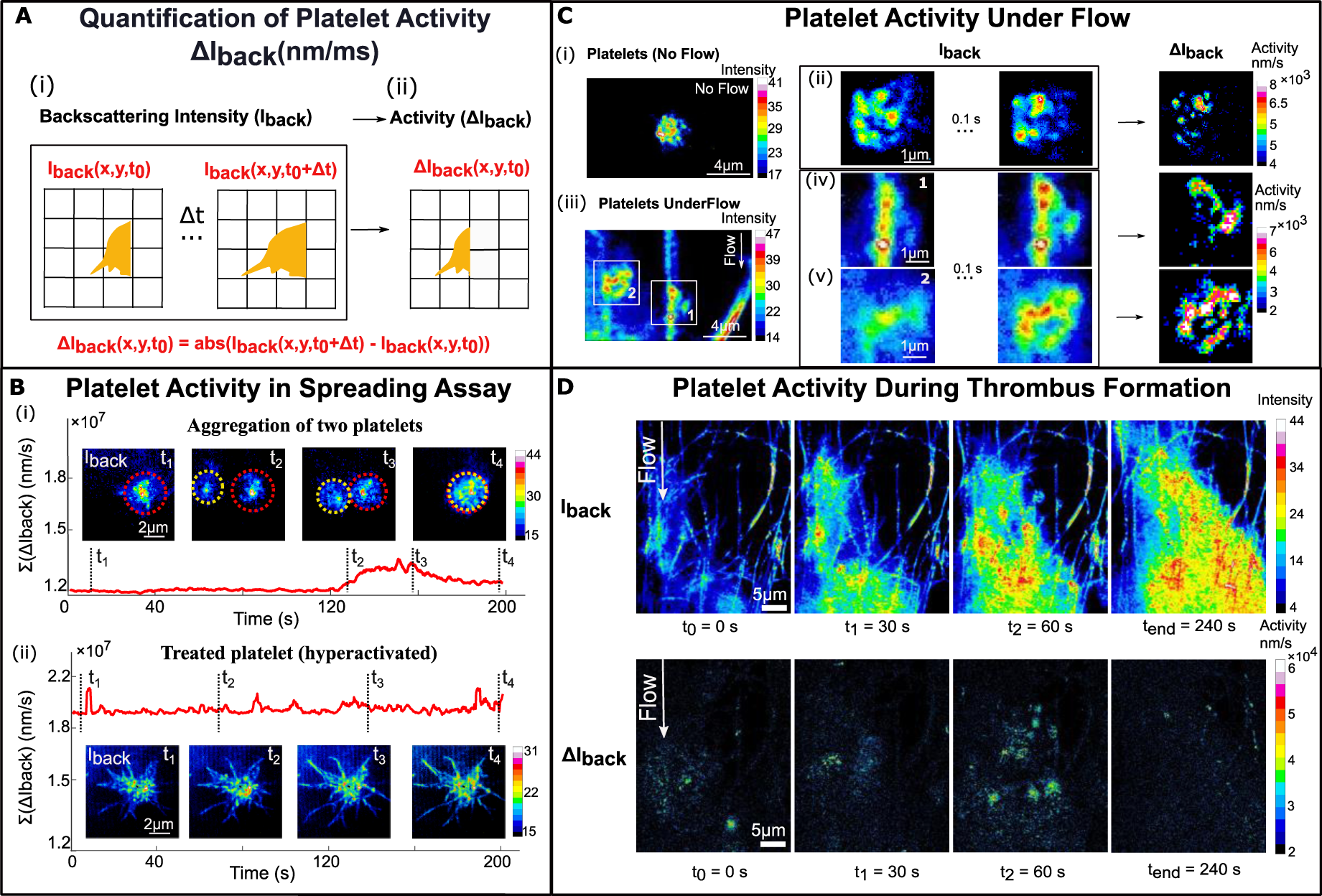
Backscattering signals of platelets in thrombi formed over collagen coated-microfluidic channels under flow. (**A**) Method used to quantify rate of change in dynamic events. Grid represents individual pixels on the camera. (i) The intensity distribution I (x,y,t_0_) of the platelet at t_0_ and after time interval Δt due to change in refractive index between the object and the medium resulting from lateral and axial position shift. (ii) Activity (rate of change) is quantified as the absolute intensity distribution difference ΔI(x,y,t_0_) (**B**) Total activity variations ∑ΔI(x,y,t_0_) within a fixed field of view over time of (i) two platelets interact and aggregate and (ii) single platelet treated with 10 μM calcium ionophore. The activity measurements shown here is the summation of activity values across all pixels in the field of view and the curve is plotted with moving average of 10 frame. (**C**) Single platelet activity quantification (i) Single platelet spreading on a fibrin-coated coverslip (under no flow). (ii) Two images of the spreading platelet taken 30 ms apart with the subtracted intensity difference shown (right). (iii) Scattering signal of single platelets under flow in a microfluidics channel where platelets move along the collagen fiber in ROI 1 or adhere to a collagen fiber in ROI 2. Two images in ROI 1 (iv) and ROI 2 (v) taken 100 ms apart with the subtracted intensity difference shown (right). **(D)** The backscattering signal at 0 s, 30 s, 60 s, 120 s and 240 s illustrate thrombus formation in a time series and the corresponding platelet activities calculated using the subtraction method (see Methods).

To test the sensitivity of this technique, i.e. level of physical motility around discrete platelets, we imaged single adhesive platelets after 20 minutes of spreading at 0.2 second time intervals with resolution of 40 nm/pixel <u>(full Movies are available in S3 and S4)</u>. Fig. 3B shows time series images of two platelets that begin to not only interact but also aggregate (t_1_ to t_4_). Here, it is clearly observed that the magnitude of the platelet activity only increases upon coming into contact with a second platelet at t_2_. However, once the two platelets completely aggregated at t4, the measured activity reduces. To further validate that the magnitude of the activity, we conducted imaged a Ca^+^ ionophore treated platelet that is known to hyperactivated (33). Fig. 3B (ii) shows the time series of the measured activity of the treated platelet where there are several sudden spikes that is directly attributed with rapid extension of filopodia and protrusions. This provide clear evidence that platelet activity using oblique illuminated light scattering has high spatial-temporal sensitive that could be suitable to quantify platelet activity during thrombus formation. Next, we carried out a microfluidic thrombosis experiment so as to test the ability of COSI to capture real time imaging of single platelet recruitment in a developing thrombus with changing mass, which to-date has remained elusive.

### Real-time observation of discrete platelet recruitment into a developing thrombus under flow

At the chosen imaging depth, the scattering component of COSI permits direct imaging of the precise location of individual collagen fibers and platelets. Hence, it is then possible to analyze single platelet spreading dynamics during thrombus formation under physiological flow rates. Here, we quantified the time-dependent spatial-temporal interaction of single platelet under different flow condition at 0.1 second time intervals with spatial resolution for each pixel to be either 40 nm or 100 nm based on the optical magnification of either 263 x or 61 x. So for a single pixel shift, the spatiotemporal resolution is 40 nm/0.1 s = 400 nm/s or 100 nm/0.1 s = 1000 nm/s. Fig. 3C shows the spatiotemporal quantification method applied to a single platelet with and without application of 1800 s^-1^ fluid shear stress (representing arterial (patho)physiological shear). For comparison, we shown spreading on a fibrin-coated coverslip without fluid flow and collagen-coated coverslip under flow as shown in Fig. 3C (i) and (ii). Using the subtraction method (subtraction method at time intervals of 0.1 s), we shown that evidence that under flow, the spatial temporal distribution of the platelet activity increases under flow. Under static conditions on immobilized fibrin (Fig. 3C (ii)), we observe a lower number of spatiotemporal interactions (averaged platelet activity 480 nm/s). However, under flow in Fig. 3C (iii)-(v), two platelets are observed to physically attach onto collagen fiber bundles under fluid shear respectively <u>(full Movies are available in S5)</u>. More importantly, Fig. 3C (iii)-(v) shows the application of the image. The two adhesive platelets (Fig. 3C(iv) and (v)) is observed to have significantly increased averaged platelet activity of 1670 nm/s and 3540 nm/s respectively. The platelet in Fig. 3C (v) with a higher level of activation (almost twice as much activity) was observed to assimilating into the platelet aggregate before stability. The increased magnitude of activity spreads before being assimilate into the base appears to enable the platelet to withstand high fluid shear stress. The recruitment and activity of single platelets captured at a sub-micron level during thrombus formation under flow, demonstrates that the scattering light images contain rich real-time morphological information, not previously observed and warrants further investigation in combination with oblique fluorescence microscopy so as to correlate with actin-rich region. However, oblique illumination widefield fluorescence microscopy is prone to accepting out of focus fluorescence that degrade image quality (34) which can be elevated using structured illumination microscopy (34, 35).

Next, we move onto measuring spatiotemporal dynamics to an entire developing thrombus (Fig. 3D) using coated collagen layer. As with the single platelet images in Fig.3C, the collagen bundles act as a fiducial marker to focus the imaging field of view and site of activation. This ensures that we capture early adhesive platelet events before they assimilate into a platelet aggregate. We adjusted the incident angle of the illumination to 50° to achieve a backscattering imaging depth of field of just around a single platelet thick (z = 4 µm). The spatial temporal map (Fig. 3D) captures the surface activity of platelets accumulating into a thrombus at 1800 s^-1^ shear rate in a microfluidic channel, based on the monotonically increasing scattering intensity from 0 s, 30 s, 60 s, 120 s and 240 s <u>(full Movies in S6)</u>. Using the subtraction method (Fig. 3D), we observed that the magnitude of activity did not increase at all that is indicative of a highly stable basal surface during thrombus formation which is consistent with existing thrombus formation. The full events of platelet spreading and interacting with collagen fiber bundles under fluid shear are shown in Movies S2 and S3.

### Mapping intra-thrombus stability and adhesive platelet activity in

So far existing high resolution platelet imaging (3, 36, 37), that has not been able to quantify platelet-platelet activities in entire thrombus and even less is known about the overall impact on the thrombus mass (38). To demonstrate the strength of combinatory label-free imaging of COSI, we investigate how COSI can extract both the intra-thrombus stability and platelet activity in relation to thrombus embolization (loss of platelet aggregates from the thrombus) as illustrated in Fig. 4A. More importantly, we aim to use COSI to quantify distinctive stages of thrombus formation. Fig. 4B-E shows multiparameter quantification capacity of COSI permitting simultaneous measurement of thrombus mass and surface activities during embolization under (patho)physiological fluid shear. Overall thrombus events were determined by spatially mapping out changes in mass and optical scattering signals of the thrombus across each imaging frame <u>(full Movies in S7 and S8)</u>. Fig. 4B (i) and depict two reconstructed QPM images of thrombi formed under 7500 s^-1^ shear stress, taken 5 minutes apart. A loss in thrombus mass due to platelet detachment was observed and the highly activated adherent platelets (also presumably procoagulant), displayed strong scattering signals between imaging frames ∼ 30 ms apart (Fig. 4C (i)). Interestingly, after applying the subtraction method we observed significant spatial-temporal surface activities of the adherent platelets (z = 4 μm). To describe mass and adhesive platelet activity variation during embolization event, a 2D map of mean mass change (Δ*m*) and standard deviation of membrane activity signal (ΔI_back_) at a shear rate of 7500 s^-1^ taken 5 minutes apart, are plotted and shown in Fig. 4D and 4E. From the surface activity maps, we observed a highly heterogenous distribution of gain and loss of thrombi mass across the 5 minutes of high shear rate exposure. Interestingly, the adherent platelet activity was spread in a concentric pattern with cores of high activity surrounded by regions of lower activity (Fig. 4D and 4E). Full data set of 2D maps of mass and platelet activity of three donors is shown in Fig. S3.

**Fig. 4.**
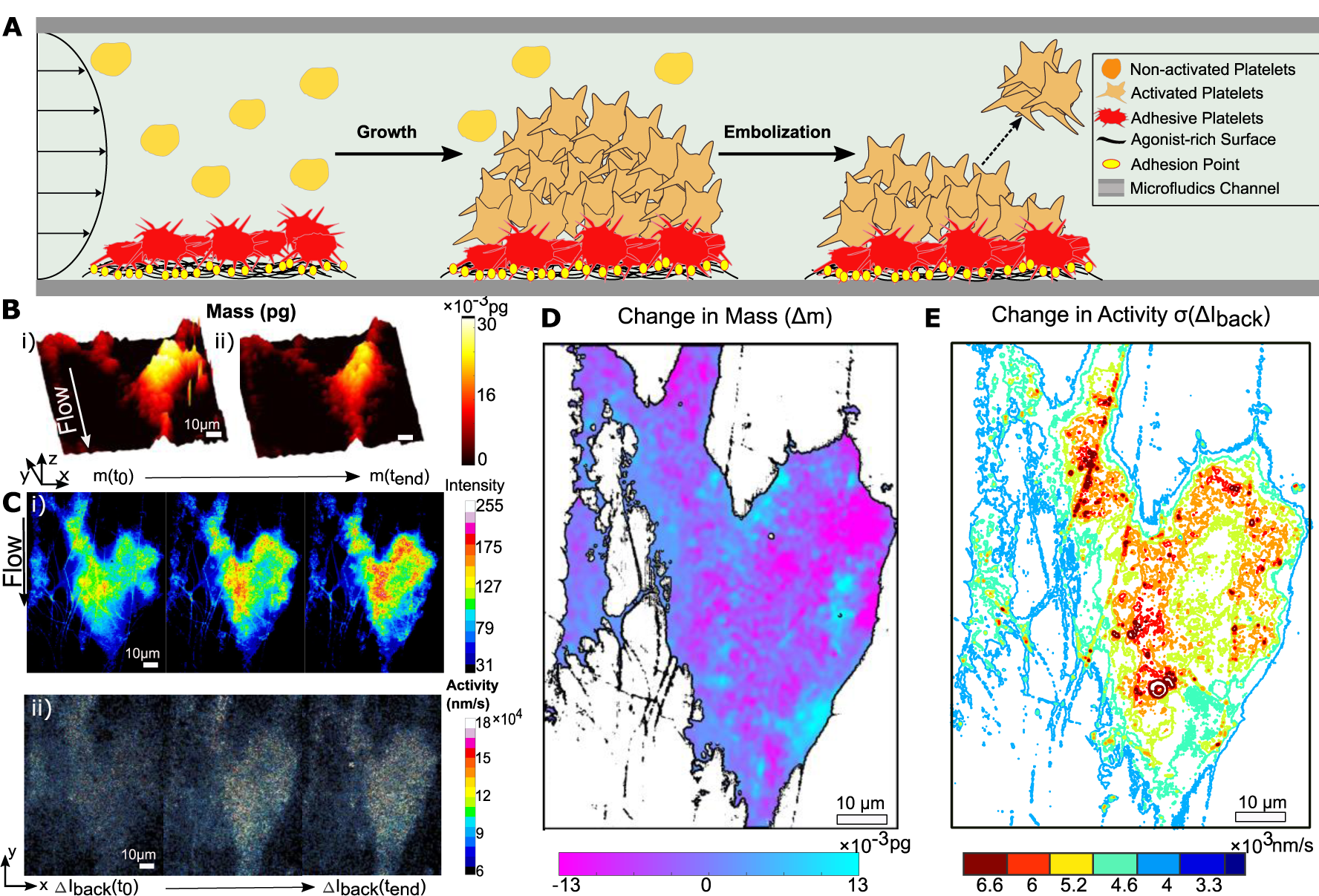
Quantifying intra-thrombus stability (changes in mass) and adhesive platelet activity during embolization. **(A)** Schematic of thrombus stability (growth and embolism) tested by the experiment procedures. **(B)** The mass change of the thrombus during embolization is defined as the mass difference between a given time from (i) start (***m***(***t***_***0***_)) to (ii) the end (***m***(***t***_***end***_)). **(C)** (i) Back scattering signal and (ii) resulted subtraction plot showing surface activity at 30 ms of the thrombus during embolization. The signal in subtraction plots is magnified 10 times for contrast enhancement. **(D)** Example of 2D plot showing change in mass distribution of a thrombus exposed for 5 min to 7500 s^-1^ shear stress with purple representing mass loss and blue representing mass gain. The region and border line of the thrombus (black line) is determined by thresholding and tracing using an activity map. **(E)** Example of a corresponding 2D activity map. The activity map is calculated with the standard deviation of activity across the imaging session and plotted using a contour plot where red and blue represent more and less activity respectively. Scale bar: 10 μm.

### Investigating the role of receptor shedding on thrombus mass distribution during formation and embolization

In order to evaluate spatial temporal behavior between mass variation and the level of adhesive platelet activity at different stages of thrombus embolization shown in Figure 4 for different donors, we plotted the change in mass *versus* platelet activity on 2D scatter plots in Fig. 5A at 30 second intervals over 2.5 minutes at a shear rate of 7500 s^-1^ (full data set at 5 minutes is shown in Fig. S4). Using the scatter plots, we observed that, after the first 30 seconds, the scattered points are more dispersed horizontally (9.5 % increase in standard deviation), reflecting higher platelet activity (Fig. 5A). At 30-120 seconds, the activity gradually increases while the mass exchange remains consistent, with minimal change in activity and mass distribution observed after 120 seconds. This suggests a level of thrombus stabilization has been achieved.

**Fig. 5.**
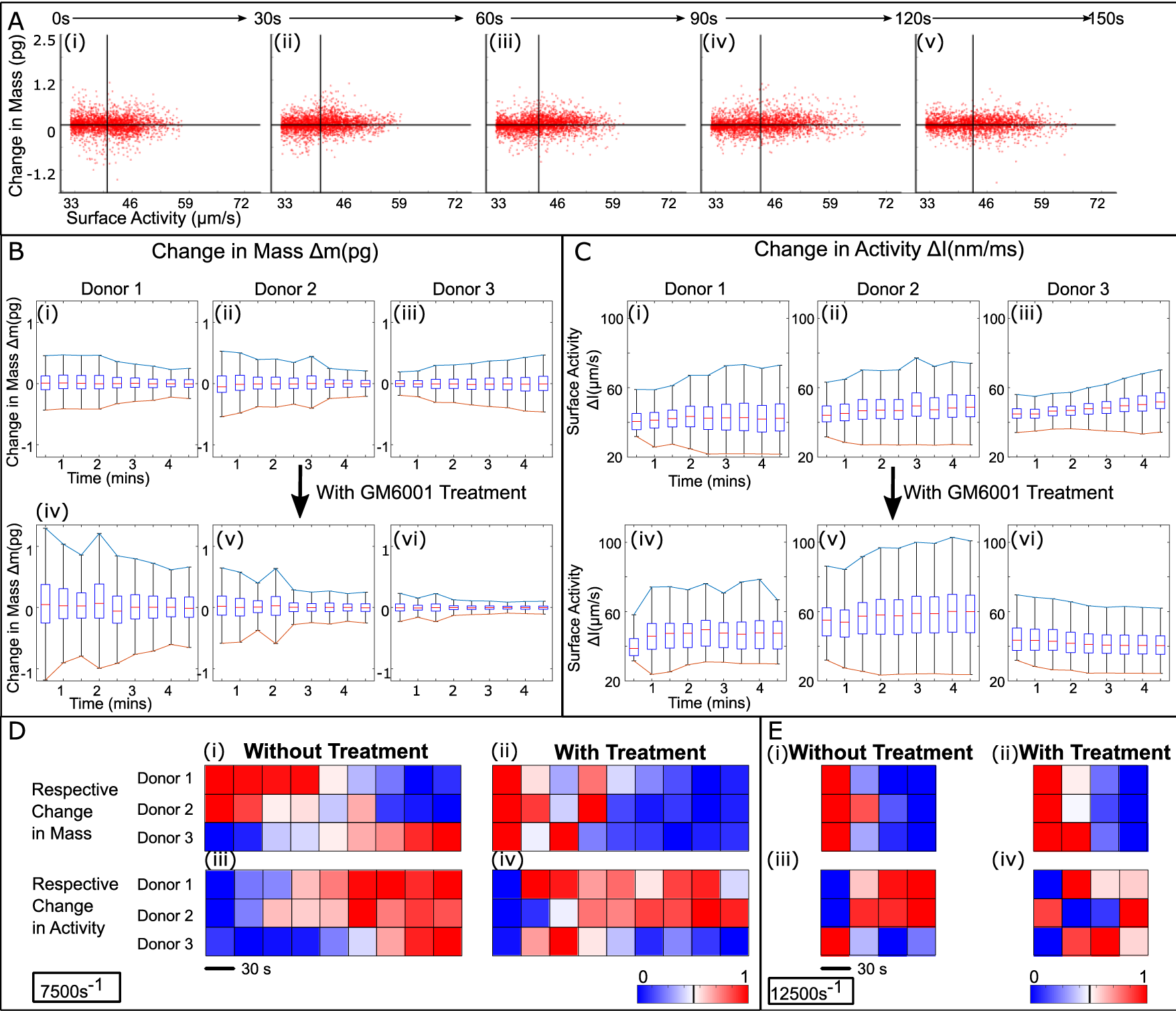
Mass and activity changes in thrombi of multiple donors with and without metalloproteinase inhibition (GM6001 250 μM; broad metalloproteinase inhibitor). **(A)** Scatter plot of thrombus under 7500 s^-1^ shear rate at 30 s time interval showing the corresponding changes in mass and activity of individual binned pixels after spatially mapping the 2D activity and mass map of the thrombus with a binning size of 10*10 pixels. The scattered plot is divided into four quadrants distinguishing loss and gain in mass and activity level. Boxplots from scatter plot are used to represent the overall trends of **(B)** mass and **(C)** platelet activity change over time. The upper and lower interquartile limit of the boxplots are drawn to highlight the interquartile range. Color coded plot of respective change in mass and surface activity of the developing thrombus under (**D**) 7500 s^-1^ and (**E**) 12500 s^-1^ shear rates with and without 250 μM GM6001 pre-treatment. The respective change is calculated as the ± 1.5 interquartile range of each boxplot.

Upon activation, platelets shed the ligand-binding ectodomain portion of key adheso-signalling receptors (39). We investigated whether prevention of activation-induced metalloproteolytic cleavage of receptors such as glycoprotein (GP) Ib-IX-V complex (40) and GPVI (41, 42) which are important for thrombus stability, resulted in changes to surface activities (as measured using COSI). To evaluate whether a more stable thrombus is formed under conditions where proteolytic receptor shedding was blocked, PRP, form three different donors was pretreated with 250 µM GM6001 (a broad range metalloproteinase inhibitor) or vehicle control, and mass exchange and surface activity in time series were compared. Donor-to-donor variation in mass change and surface activity of thrombi imaged was observed (Fig. 5B and C (i-iii)) in vehicle or GM6001-treated samples. Although heterogenous thrombi behaviors were observed between different donors, in PRP treated with GM6001, there was a consistent tendency for thrombus mass to decline with time, indicating reduced mass exchange and a more stable thrombus when receptor proteolysis was blocked (Fig. 5B iv-vi). However, the basal platelet activity showed generally little change with GM6001 treatment (Fig. 5C iv-vi). Fig 5D and 5E show color coded plots that represent the respective change in mass and activity (based on normalization of the ± 1.5 interquartile range), with and without GM6001 treatment of thrombi exposed to 7500 s^-1^ and 12500 s^-1^ shear rates. Again, mass exchange was observed of all donors at both 7500 s^-1^ and 12500 s^-1^ shear rates with GM6001 treatment, which reached a steady state after the distinct drop over time. The surface activity, on the other hand, shows more variation between donors. We observed lower increment (donor 1 and 2) or a slightly decreased (donor 3) trend at 7500 s^-1^ and activity at 12500 s^-1^ shear rates after treatment. This reduced or minimized trend in surface activity is consistent with the basal surface becoming more stable, corresponding with minimum mass exchange.

## DISCUSSION

Platelet spreading assays have previously been studied with label-free imaging modalities including QPM (43), brightfield microscopy, RICM (19, 44) and even with scanning ion conductive microscopy (45). Among these techniques, brightfield and RICM can only provide simple morphological information for single platelet and not readily used to quantify single platelet-level activity during thrombus formation. Additionally, RICM makes use of near-field effect that is only sensitive to a depth of field of ∼100 nm above the coverslip as contrast is generated from the phase shift between reflected light from the coverslip and scattered light from the sample. QPM and scanning ion conductive microscopy can provide topographic information but fails to measure the adhesion site between platelet and prothrombotic surface. In this paper, we conclusively demonstrated that COSI achieves high contrast imaging single platelet that can readily quantify the magnitude of platelet activity measured at the adhesion site together with quantitative mass measurement during platelet spreading assays that will be crucial to unpacking complex biochemical and bio-rheological properties.

Here we shown that COSI can achieve real-time correlative quantification of both thrombus mass and thrombus surface activity on coated microfluidic chips and so provide a new means to quantify platelet activities under flow. This is especially because COSI imaging is a multiparameter tool that can unpick the heterogenous distribution of different thrombus interacting with different types of substrate-coated surfaces. These types of multiparameter imaging measurements will advance our understanding of how platelets interact with various agonists, for example to assess differences between GPVI interactions with collagen, fibrin and other GPVI binding proteins, and the effects the different interactions have on thrombus stability, which is important for future anti-thrombotic therapeutic development. In addition, we expect combinatory approaches of quantitative label-free microfluidic imaging tools open up new clinically relevant opportunities to evaluate different platelet subpopulations in relation to the thrombus structure and how they function during thrombus formation under fluid shear force.

Since the main signal here is refractive index change, it is likely that the activities measured at the platelet-substrate interface reflect cytoskeletal rearrangement (actin refractive index is 1.57 (46)), due to similar actin arrangement observed along membrane ruffling when using phalloidin staining and fluorescence microscopy (Fig. S2C and S2D). Although the scattering signal from inside of the platelet (Fig. S2E) does not corelate with the fluorescence signal (Fig. S2F), which might due to a contribution from degranulation to light scattering (47). Future studies using inhibitors of cytoskeletal rearrangement, signaling or degranulation will better describe the circumstances of this newly recognized light scattering signal from single platelet.

The COSI imaging approach could also help address where procoagulant platelets, characterized as having amplified surface area, sustained calcium rises and phosphatidylserine (PS) exposure are located in newly formed or embolizing thrombi. Procoagulant platelets are thought to be highly motile in order to accelerate coagulation (48) for forming fibrin-rich hemostatic plugs, that form within a thrombus and are mechanically translocated to the periphery via thrombus contraction (49). As COSI and other refractive index-based imaging tools do not require uptake of membrane-permeable dyes, or fluorophore-conjugated antibodies, which can disrupt function of important platelet receptors or surface molecules (50), these tools are ideal to be developed for utility in clinical and drug evaluation settings.

COSI conducted quantitative imaging under two levels of fluid shear stress and identified interaction of clusters of platelets with collagen fibers at high spatial temporal dynamics at nanometer per second resolution. This enabled real-time longitudinal temporal evaluation of the impact of metalloproteinase inhibition on platelet aggregate formation and stability that has not been done before. The correlation of the mass exchange and change in surface activity of the developing thrombus indicated the significance of intra-thrombus mass exchange at picogram levels and stabilized surface activity after treatment of a broad range metalloproteinase inhibitor. The COSI system therefore can easily be expanded to test the effect of new and existing anti-thrombotic agents and specific receptor antagonists on thrombus size and stability in future studies. Further COSI approaches could be expanded for use in clinical situations, to help identify platelet defects in patients with unexplained bleeding or diagnosing embolism risk.

## Materials and Methods

### COSI Microscopy Platform

The COSI microscopy platform setup is illustrated in Fig. S1. An inverted microscopy imaging platform was chosen to construct COSI using two separate beams of different wavelength; a 632.8 nm HeNe laser (HeNe laser, TEM_00_ mode, JDS Uniphase, 1144p-3581) for off axis QPM and a second linearly polarized 488 nm laser (Stradus 488-150) for rotational backscattering illumination. To co-locate the interferometric and backscattered signals, two illuminating beams share the same microscope objective (Olympus, 60X, NA 1.49, oil immersion). To ensure both illumination beams reach the sample as close to ideal plane waves, the beams are focused onto the back focal plane (BFP) of the respective objectives; transmission objective (10X, air) for interferometry and oblique illumination objective (Olympus, 60X, NA 1.49, oil immersion). Two cameras were used to collect the transmitted interferogram (7) and summed backscattering signal (22), respectively.

On the transmitted light path, the beam is first split into two arms (object and reference) by a standard dual fiber splitter. The objective arm illuminates the sample and recombined with the reference by a 50/50 beamsplitter (Thorlabs, BS013) to form the interferometric pattern that is digitally recorded with a sCMOS camera (Thorlabs Quantalux CS2100M-USB).

On the rotational backscattering imaging path, the beam is controlled using a 2 – axis galvanometer mirror (Thorlabs, GVS212/M) that conjugates to the back focal plane of the imaging objective (Olympus, 60X, 1,49, oil immersion). The scanning rate of an orthogonal pair of galvanometer mirrors fixed at 3600 Hz (24 scans per cycle) to be synchronized with the shutter of the camera of 150 Hz, which results in a maximum imaging speed of 150 fps. This synchronization allows a high speed 2D complementary metal oxide semiconductor (CMOS) to capture one full cycle of rotational scanning and sum it directly. To remove the back reflection of the illumination beam and only allow backscattering to reach the CMOS (Point Grey Flea3 USB3, FL3-U3-13Y3) imaging detector, a diaphragm (Thorlabs, ID15/M) is placed at the Fourier plane of the imaging lens. The total imaging depth of view can be tuned from 200 nm to 4.3 µm by applying different oblique illumination angles (90° to 45°). In doing so, the amount of scattering light imaging from a given imaging depth can be customized as required.

Two different imaging fields of view with magnifications of 263 x and 61 x are used depending on the pixel resolution required for each experiment, calibrated using a 1951 USAF target (Edmund Optics, #38-256) before each experiment.

### Pixel Mismatch Adjustment

The two digital cameras used for scattering and interferogram possess different pixel resolution resulted in a mismatch of 0.05%. This was removed using digital scaling of the image using the USAF1951 calibration grid.

### Scattering Signal and Refractive Index Calibration

The backscattering is generated when the illumination encounters a refractive index difference between the sample and medium as we illustrated in Fig. S1A (i) and 5.2 μm microspheres were used to illustrate this effect as shown in Fig. S1A (ii). Multiple backscattering signals results in the highest signal to noise ratio along the perimeter of the microsphere that appears to coincident with largest refractive index differences between the interface of the cover glass and the microsphere. Additionally, COSI, through the magnitude of backscattered intensity, also quantitates the refractive index difference of the microspheres and medium. Fig. S1B shows the COSI scattering signal with a mixture of microspheres with average refractive index differences ranging from 0.035 to 0.25. The measured scattering signal (signal to noise ratio – (SNR)) was plotted against the refractive index difference. The calibration between refractive index difference and scattering signal is done using microspheres made with polystyrene (3 µm diameter and refractive index 1.59) and silica (5.2 µm diameter and refractive index 1.51) embedded in sucrose solutions with varying refractive index ranging from 1.334 to 1.45 (HI 96802 Hanna Instruments). To ensure consistency, the incident angle of the beam to the objective is altered to ensure the sample is illuminated under same oblique angle (50°). In order to quantify signal to noise ratio, we define the scattering intensity as signal and imaging background as noise in our system.

To calibrate the stability of the SNR measurement through COSI system, we quantified the SNR variations from continuous backscattering signal of different samples under different flow conditions as shown in Fig. S5. Fixed microsphere (5.4 μm) with no flow and coated collagen fiber in microfluidics chamber are shown in Fig. S5A-B and C-D respectively. We plotted SNR over imaging time across three field views for both samples and we did not observe significant change in SNR throughout imaging period as shown in Fig. S5C and D.

Based on the SNR measurement of microsphere in different medium, we calculated the SNR of scattering signal from a single platelet membrane ranged from 1.19 – 1.33. Using the silica bead calibration plot (Fig. S1B), we obtained an estimated refractive index of 1.40 that agrees with measurement of n_platelet_ ≈ 1.399 (51) with the assumption that n_pbs_ ≈ 1.335 (52).

### Depth of Field Calibration

The imaging depth of field of scattering light was tailored by changing the oblique illumination angle. In order to visualize different depth, we suspended a mixture of microspheres of 200 nm and 5.2 µm polymer sphere under two oblique angle ∼ 90° (close to total internal reflection) and 50° as shown in Fig. S1B (i) and (ii) and the corresponding scattering images in Fig. S1B (iii) and (iv).

The imaging depth was then calibrated by moving a single layer of 1 μm microspheres along the axial direction at 1 μm increments. Scattering images were collected and plotted in Fig 2F, showing that for each illumination angle, the normalized mean intensity distribution along the image. The imaging depth is estimated by evaluating the full width half maximum of the fitted Gaussian curve (goodness of fit ∼ 0.95) as shown in Eq.1.

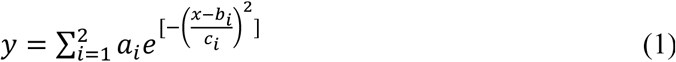

### Phase Retrieved Mass using Off-axis Interferometry

Off-axis quantitative phase microscopy (QPM) is well-suited to measure dry mass of a developing thrombus without labeling (53). A change in refractive index provides a signature optical phase delay that can be recorded and quantified through interferometry and optical scattering (54). The optical phase difference is measured between two coherent light paths; the transmitted light path through the sample and another projected directly onto the camera. The recorded interference intensity pattern is then numerically processed to retrieve the phase delay *ϕ*_*t*_(*x,y*) (54, 55). For a given refractive index difference (Δn) (51), a phase delay *ϕ*_*t*_(*x,y*) is imposed onto the transmitted light for a narrowband wavelength light source. This refractive index change of biomolecules approximate change in dry mass and can be determined from changes in phase (26, 27, 53, 56).

The mass of a homogenous sample can be directly calculated using volume and density (d*v). The mass of the polystyrene microsphere is calculated with d = 1.05 g/cm^2^ (26, 54). For biological cells of a given refractive index, the linear increases in dry biomass (proteins and nucleic acids without water) falls within a very narrow range of approximately 2 × 10^−4^ m^3^/kg that allows one to estimate the dry mass of thrombi and cells (26, 53). The equation used to reconstruct dry mass of thrombi from phase *ϕ*_*t*_(*x,y*) is shown in Eq. 2, where α is the specific refractive increment. Change in dry mass of the thrombus identifies the affinity/recruitment of platelets into aggregate which is determined by the mass difference before and after the flow of saline (0.9% NaCl) as shown in Eq. 3.

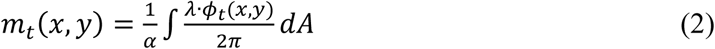

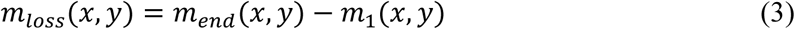

### Platelet Activity calculation from backscattering signal

An activity map registers platelet activity or movement; depending on the rate between frames used for each subtraction as shown in Eq.3. In COSI, the standard deviation of the activity is calculated so as to identity object movement between imaging frame, hence, we define this measurement as change in activity as shown in Eq. 4.

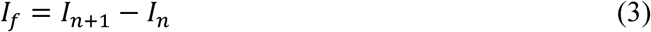

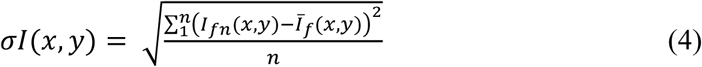

### Scatter Plot of Platelet Mass and Activity

A binary mask was generated by thresholding the 2D backscattering map to filter out the background on the mass and backscattering maps so that only the thrombus region was quantified. We binned a series of pixels (10 × 10 pixels) on the filtered mass distribution map and backscattering map, respectively. Each binned pixel was assigned the value of mass change to the y axis and the standard derivation of scattering intensity to x axis on the scatter plot to generate one scatter. The data points that fall more than 3 standard deviations from the mean were removed to show the distribution of the scatter plot. The data points were then classified into 4 different clusters by defining threshold values for mass change and scattering intensity. The threshold value for mass is 0 pg which differentiates gain and loss in mass, while the threshold for scattering intensity defines low and high level of activity and was calculated from the mean value of the intensity for all the data points.

### Platelet Preparation

Venous whole blood was collected in anticoagulants, 3.2% trisodium citrate (for platelet-rich-plasma (PRP) experiments) or acid-citrate-dextrose (ACD; 97 mM trisodium citrate, 111 mM glucose, 78 mM citric acid), for single platelet experiments. To obtain PRP, whole blood was centrifuged at 110 *g* for 20 minutes at room temperature, no brake. For metalloproteinase inhibition experiments PRP was pre-treated with 250 µM GM6001 >10 min prior to experiment commencement. To obtain washed platelets, ACD-treated PRP was centrifuged at 1271 *g* for 15 minutes at room temperature with high brake. The platelet-poor plasma (PPP) was removed and the platelet pellet was resuspended in 123 mM NaCl containing 13 mM trisodium citrate and 33.3 mM glucose, pH 7.0 and then centrifuged at 1271 *g* for a further 15 minutes. After repeating this washing step three times, the platelet pellet was resuspended in Tyrode’ s buffer (5.5 mM glucose, 137 mM NaCl, 12 mM NaHCO_3_, 1.8 mM CaCl_2_, 0.49 mM MgCl_2_.6H_2_O, 2.7 mM KCl, 0.36 mM NaH_2_PO_4_.H_2_O, pH 7.4) to reach a platelet count of 2 × 10^7^ platelets/mL.

### Platelets Spreading Assay

Glass coverslips (#1.5 thickness) were coated overnight at 4°C with 100 μg/mL collagen Reagent HORM (Takeda, Austria) or coated with 100 μg/mL human fibrinogen for 30 min, then 1 U/mL thrombin for 15 min, followed by heparin 50 U/mL for 15 min (to neutralize thrombin). The coated coverslips were rinsed with phosphate-buffered saline (PBS) then blocked for 30 min with 5 mg/mL heat-inactivated bovine serum albumin in PBS, to minimize non-specific binding. Coverslips were rinsed before addition of 50 μL of platelets (∼2 × 10^7^ platelets/mL) and morphology was monitored using COSI at indicated times.

### Imaging Thrombus Formation and Embolization Test on Collagen-coated Microfluidic Channels under Flow

Thrombus formation under flow was assessed using microfabricated straight channels (dimensions: 400 μm width, 100 μm height and 1.2 cm length), made from PDMS (polydimethylsioxane) from Sylgard 184 (Dow Corning) with 10:1 base to curing agent ratio) using soft lithography processes (Fig. 1A).. The microfluidic channel was coated with 100 μg/mL collagen Reagent HORM (Takeda, Austria) overnight at 4°C or 2 hours at 23°C. Prior to each experiment, the channel was washed with saline at 300 μL/min for 2 minutes. PRP was then drawn into the channel (opposite input used for agonist coating) with the syringe pump (syringe pump (Harvard Apparatus PHD 2000) in suction mode for 5 min at a shear rate of 7500 s^-1^ (flow rate: 300 μL/min and shear stress: 75 dyne/cm^2^) to induce thrombus formation (Fig. 4A). The reservoir was then switched to saline and thrombi were exposed to shear rates of 7500 s^-1^ for 5 minutes followed by 12,500 s^-1^ for 2 min to mimic pathophysiological shear rates and induce embolization events.

## Author Contributions

W.M.L conceived and supervised the project. Y.Z, S.J.M, Y.J.L performed the experimental work, T.X fabricated the microfluidic devices, Y.Z and T.N.X calibrated the imaging performance of COSI. W.M.L, Y.Z, S.J.M and E.E.G interpreted the quantification of COSI data and Y.Z. carried out the data analysis along with S.J.M. W.M.L, Y.Z, S.J.M and E.E.G wrote the manuscript with input from all authors.

---

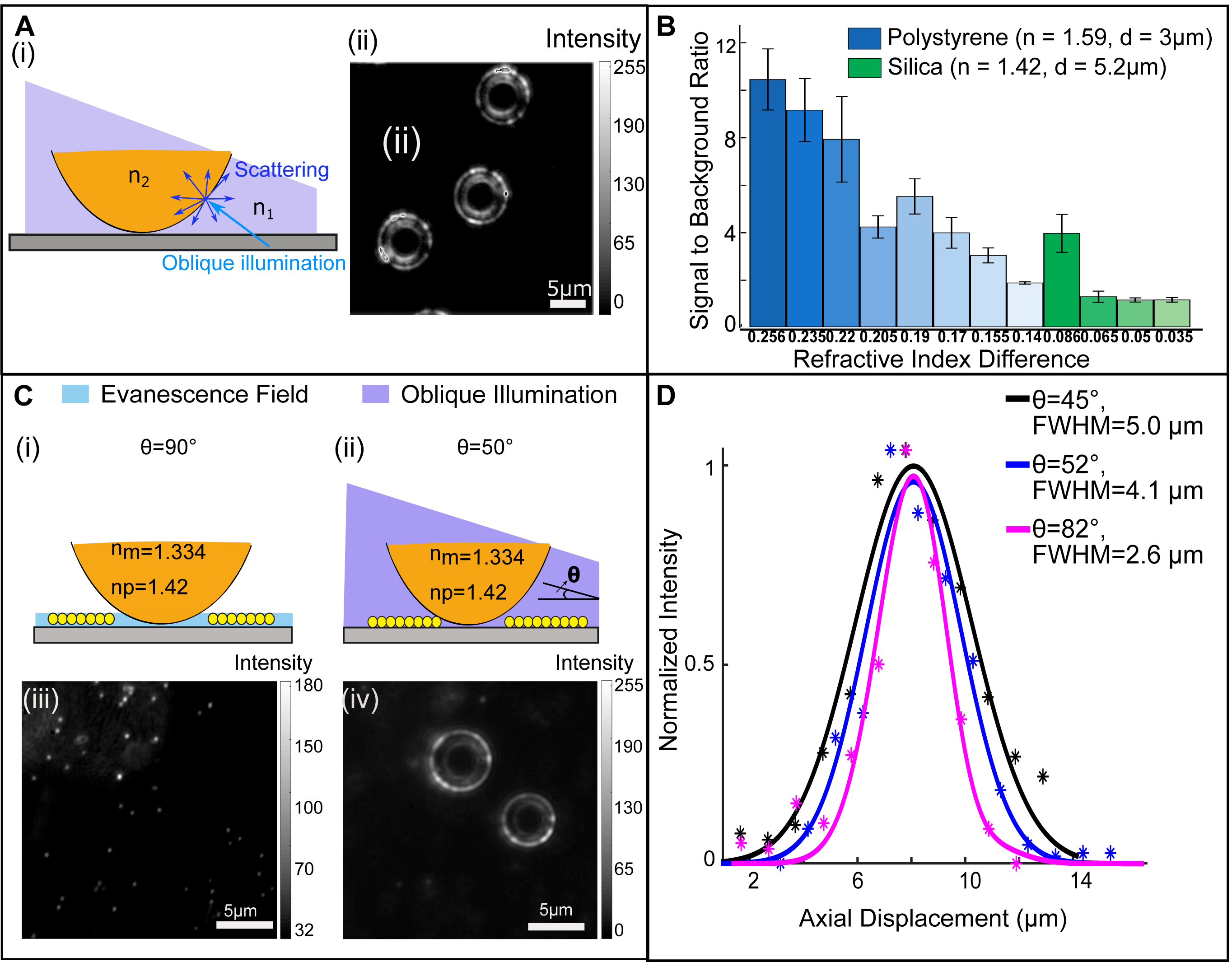

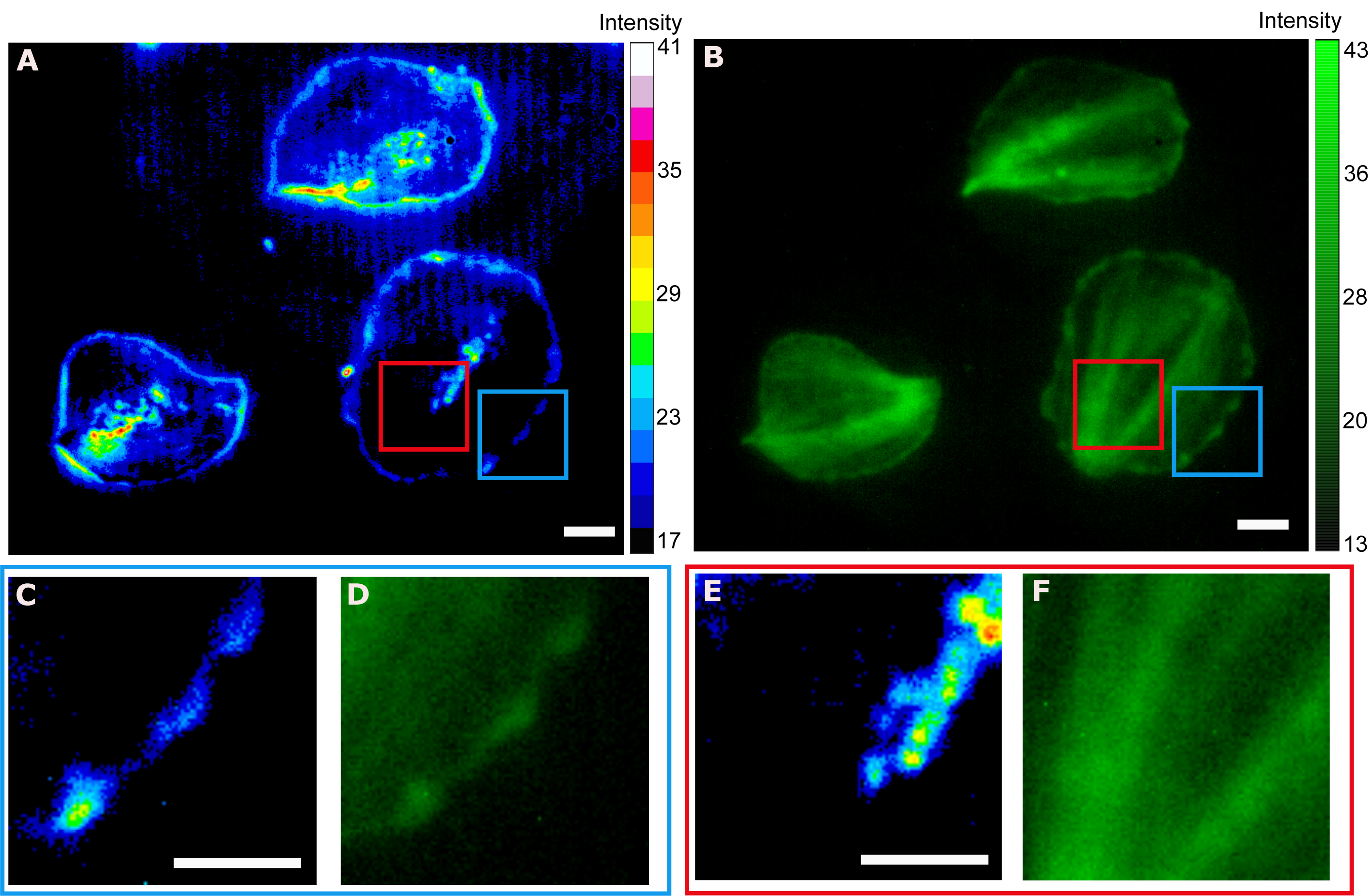

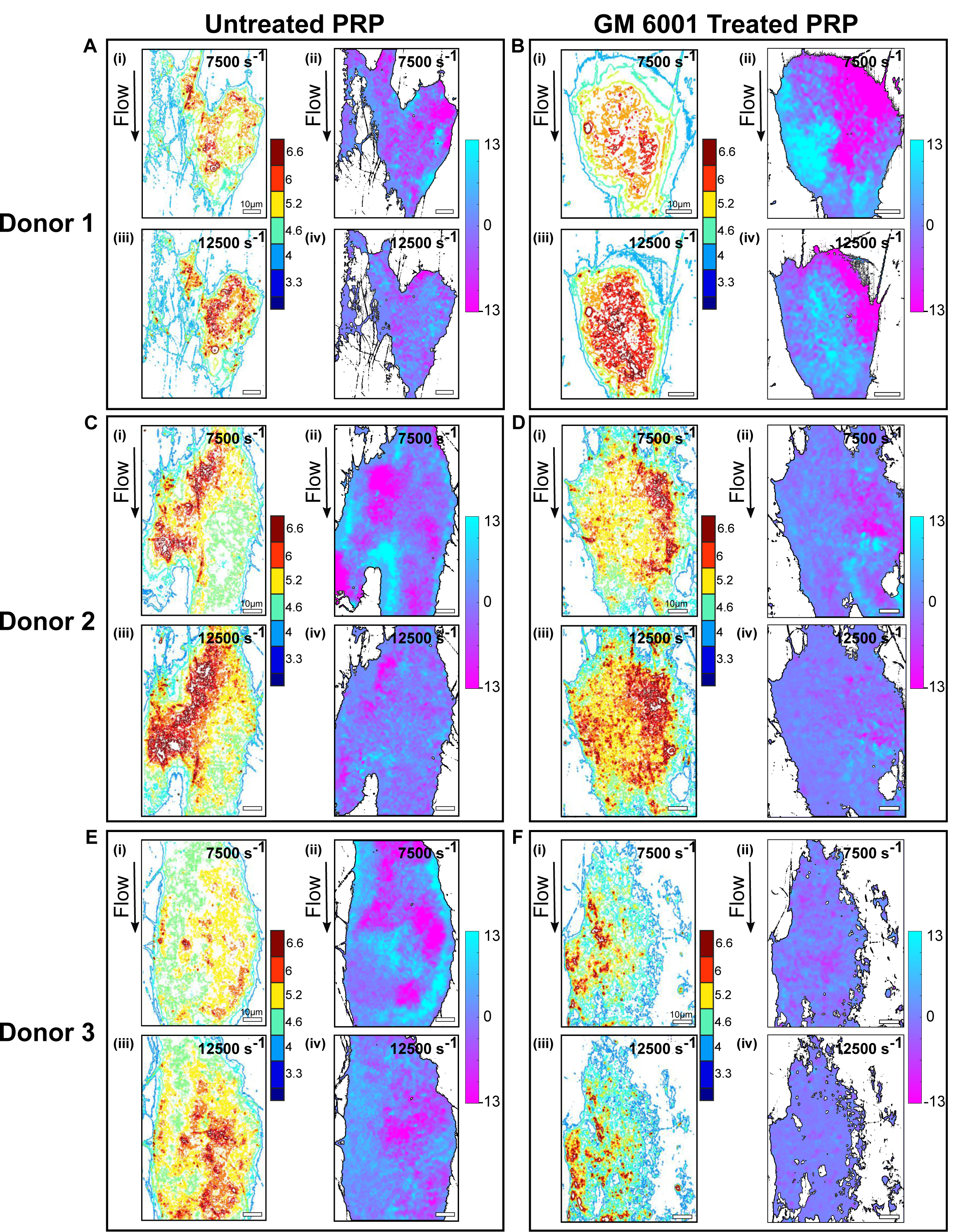

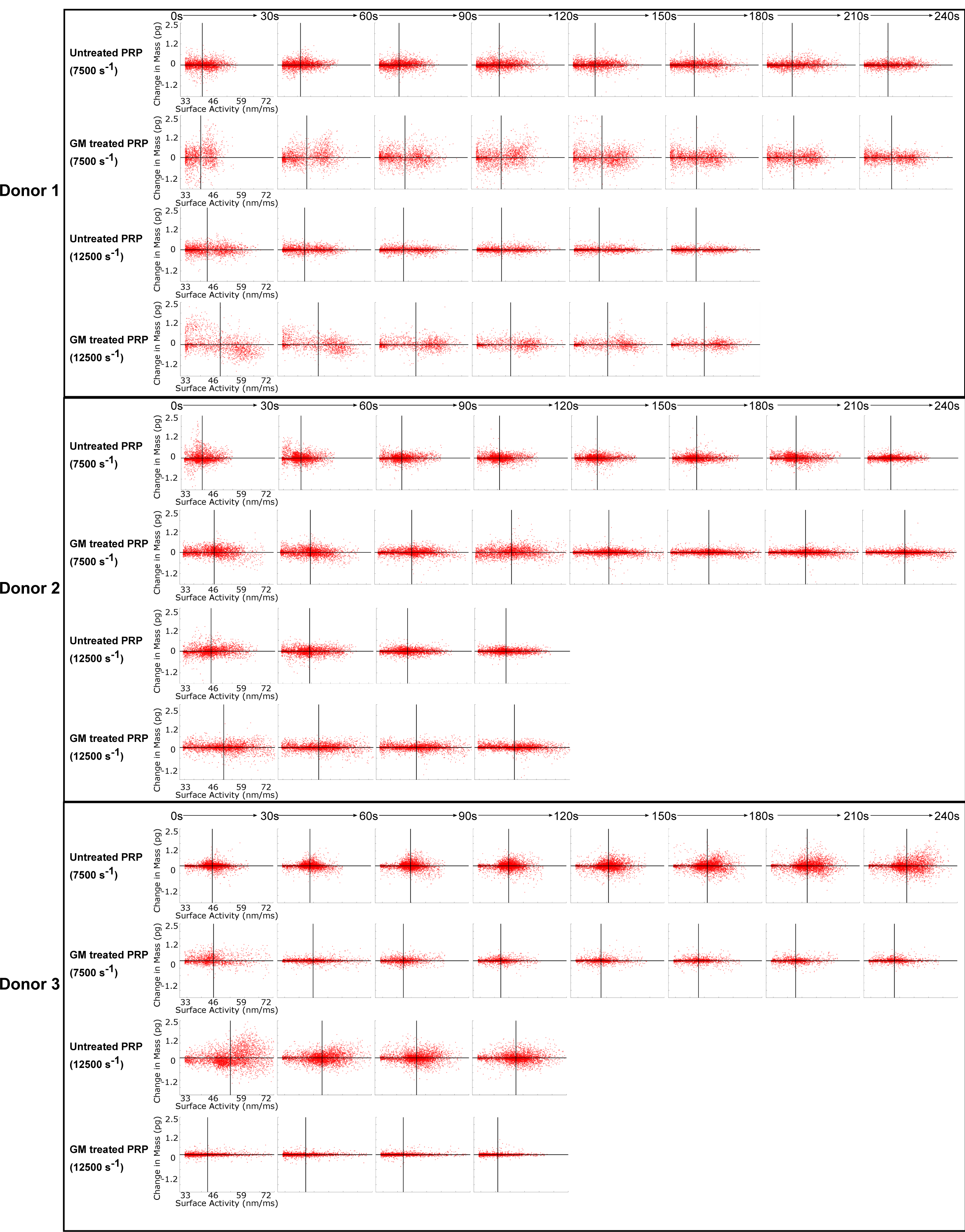

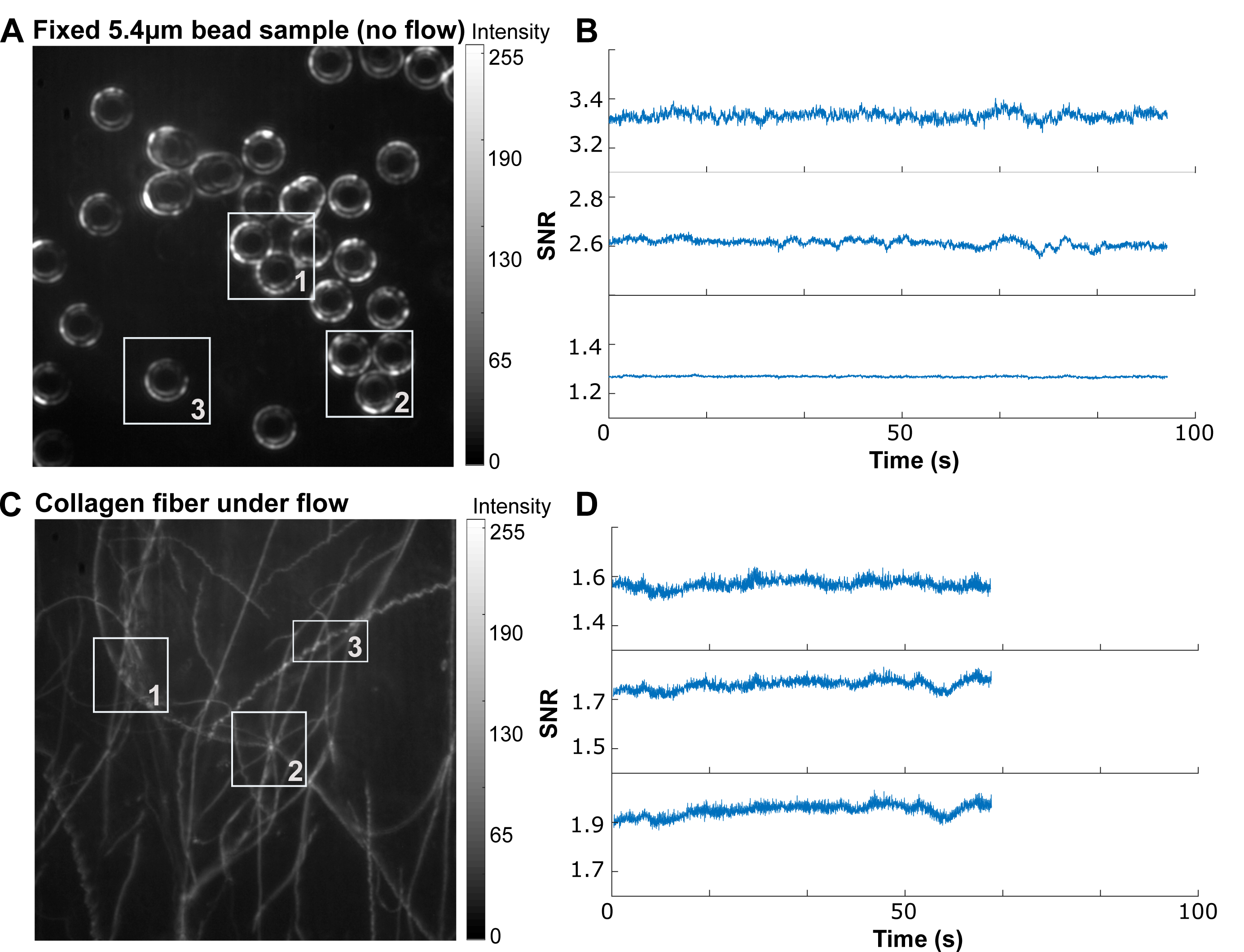

